# Sex differences in disease genetics

**DOI:** 10.1101/063651

**Authors:** William P. Gilks

**Author notes:** This manuscript is a pre-print for a review commissioned by eLS (John Wiley & Sons), http://www.els.net/WileyCDA/).

## Abstract

There is long-standing evidence for gene-by-sex interactions in disease risk, which can now be tested in genome-wide association studies with participant numbers in the hundreds of thousands. Contemporary methods start with a separate test for each sex, but simulations suggest a more powerful approach should be to use sex as an interaction term in a single test. The traits currently with the most compelling evidence for sex-dependent genetic effects are for adiposity (predictive of cardiac disease), type II diabetes, asthma and inflammatory bowel disease. Sexually dimorphic gene expression varies dynamically, by age, tissue type, and chromosome, so sex dependent genetic effects are expected for a wide range of diseases.

**Key concepts:**

1. Compelling findings of sex-dependent genetic effects on disease have been made in adiposity-related anthropometric traits, type II diabetes, and inflammatory bowel disease. Other disorders remain to be more fully investigated, regardless of what sexual differences they exhibit in prevalence and presentation.
2. Current evidence indicates that sex difference in gene expression is not required for a SNP to have a sex-dependent effect. However, sex differences in gene expression vary dynamically, by organ and age, so generalisations may be inaccurate without comprehensive data.
3. Sex-dependent risk alleles are predicted to be of greater effect size than conventional ones, because natural selection acts only against the sex which has the disease. There is evidence for this from a high-powered GWAS of adiposity-related traits.
4. Many of the large GWAS meta-analyses look for sex-dependent genetic effects by testing male and female groups separately. However, this may be under-powered compared to a whole-sample, gene-by-sex interaction test.

**Glossary:**

- **Genome-wide association study (GWAS)**. Method for identifying molecular genetic variation that controls heritable traits, in a population sample. Involves assessing the correlation between allele frequencies and phenotype value, at millions of markers of common genetic variation across the genome.
- **Sexual dimorphism**. A difference between males and females in a population for the value of a particular trait. May include anything from anatomical measurements to expression level of a gene.
- **Sex-dependent genetic effect**. A disease risk allele is termed sex-specific when it increases risk in one sex only but has no effect on the disease in the other sex. The term sex-biased is used for an allele causes a significant increase in risk of disease in both sexes, but for which the magnitude of the risk increase is significantly different between males and females. There are also reports where an allele that increases risk of a disease in one sex reduces risk of the same disease in the other sex but none have been replicated, and there is no biochemical reason why this could be true. It effectively constitutes a sexually antagonistic effect, but should be distinguished from intra-locus sexual conflict which explicitly requires than an allele have opposing effects on the evolutionary fitness of males and females (Bonduriansky and Chenoweth 2009). All of the above relationships constitute a form of sex-dependent genetic effect.

## Introduction

Knowledge of the genetic causes of disease is rapidly expanding, following recent advances in genotyping and computation methods, and massive participant numbers. There are currently two important questions. Firstly, why do results from genome-wide association studies not reach the predictions of classical quantitative genetics (Manolio et al. 2009)? Secondly, how can we use knowledge of risk genes help in the treatment of individuals (Manolio 2013)?

The modification of genetic effects by environmental variables (such as smoking, nutrient intake, stress) might help answer these questions, they are difficult to accurately quantify. Sex (gender) is comparable to an environmental variable because it influences the immediate physiological environment within which a gene functions, via the sex-determination pathway and steroid hormones. In contrast to classical environmental variables, sex is simple to measure, equally-distributed in all populations, and determined at conception. Sex-dependent genetic effects have long been known, with good evidence coming from studies of Mendelian sex-linked disorders, narrow-sense heritability, and linkage mapping of quantitative traits (Ober, Loisel, and Gilad 2008). Finally, the mitochondria, which is maternally-inherited and shows a strong sex differences in function, harbours many disease-causing mutations which effect predominantly males (Beekman, Dowling, and Aanen 2014).

There have been broad assumptions that sex-dependent genetic effects are the result of differences in sex hormone levels, but experiments using hormone treatment and gonadectomy demonstrate that sex differences in core domains of immune response, behaviour, and toxin resistance are controlled by sex chromosomes, and not by sex hormones (Penaloza et al. 2009; Ngun et al. 2011). Additionally, studies of human cell lines, which are devoid of sex hormones, indicate that expression level of up to 15% of genes is determined by the combination of both genotype and sex (Dimas et al. 2012). With genome-wide association testing and massive participant numbers, the true extent of robust and actionable sex-dependent disease associations is starting to be revealed.

Since the last review of sex-differences in disease genetics, which also provided an insight into possible their evolutionary origins (Gilks, Abbott, and Morrow 2014), many new discoveries have been made. I firstly review methods for detecting sex dependent effects in genome-wide association studies (GWAS), and then summarise the most convincing associations, and the current knowledge within disease categories. Using evidence from studies in transcriptomics, cell biology and biochemistry, I discuss the diverse mechanisms by which an allele can have a sex-dependent effect on disease risk.

**Figure 1.**
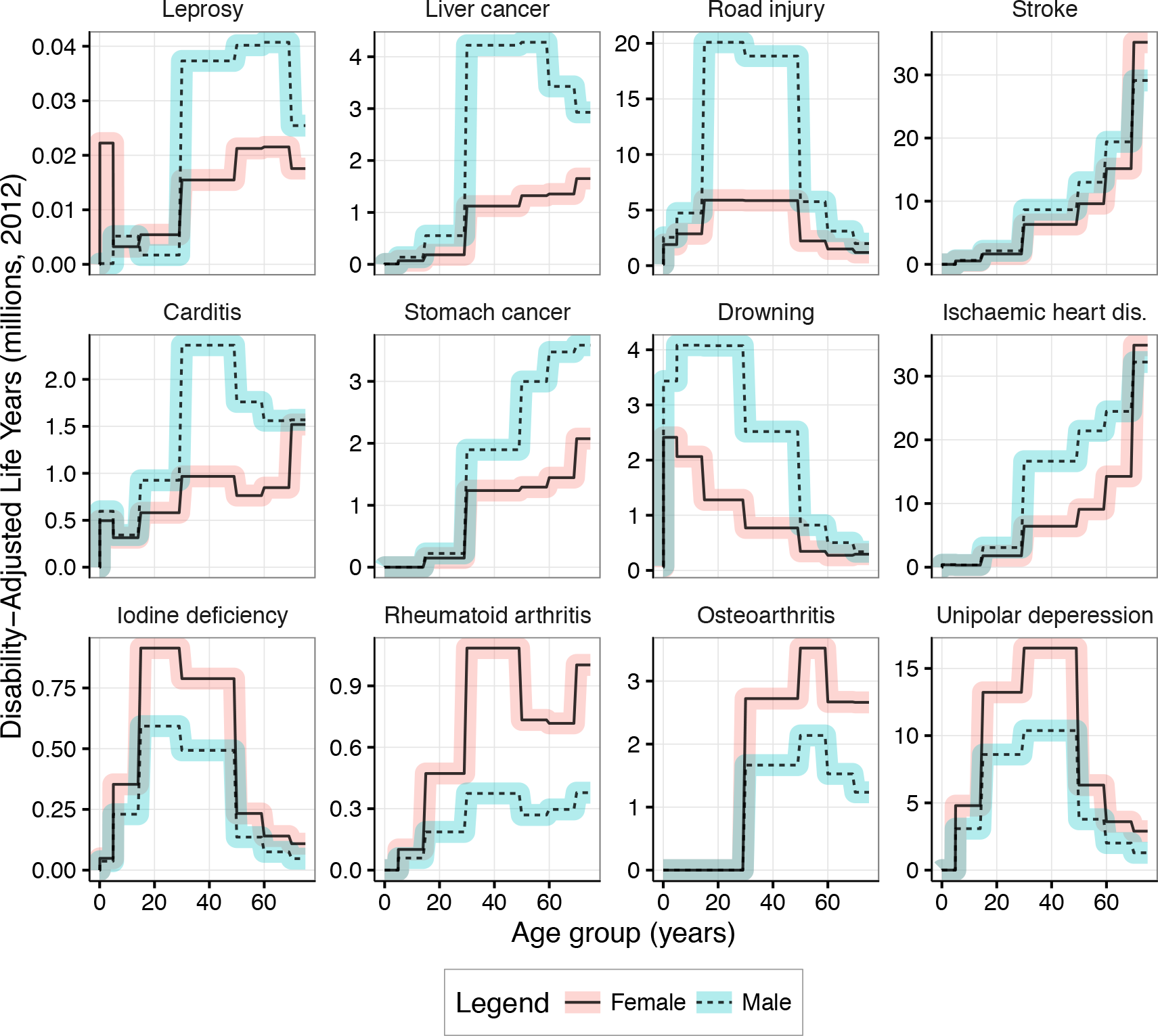
Disability-adjusted life years (DALY) by age-group and sex, for selected causes. Source data is for the global human population, year 2012 (WHO 2016). The causes were selected to show diverse and interesting patterns of sex differences. The arrangement of plots is by increasing DALY from left to right, and vertically by sex difference. Note that the y-axis scale difference between plots, so that for Stroke (top-left), the DALY is 1 million greater in middle-aged men than in women.

## Methods for identifying sex-dependent effects in genome-wide association studies

Genome-wide association studies for common diseases, typically model the relationship between genotype and disease status using logistic regression to detect additive genetic effects, whereby the effect of two alleles at a locus doubles the risk from having just one. In large GWAS with multiple sample collections, meta-analysis is appropriate. Detection of sex-dependent effects in GWAS meta-analyses has been pioneered by Magi and colleagues, with both a sex-differentiated test of association, and a test of allelic heterogeneity between the sexes, which withstood comprehensive simulations for loss-of-power (Magi, Lindgren, and Andrew P. Morris 2010). The software these authors developed, ‘GWAMA’, has been applied to many large GWAS, although without tests for sex-dependent effects.

In many recent GWAS, with sample sizes in the hundreds of thousands originating from multiple clinical research groups, t and p-values for a sex difference have been calculated from separate-sex meta-analysis statics (i.e. p, SE and *β*), where r was the Spearman rank coefficient across all SNPs (Randall et al. 2013; Winkler et al. 2015; Shungin et al. 2015)). A permutation-based variation of the separate-sex method has been described (Liu et al. 2012), but the separate-sex approach may be underpowered compared to main-effects analysis because the test statistics for each sex are from a smaller sample (Behrens et al. 2011).

The alternate method for detecting sex-dependent genetic effects in case-control data is by using sex as an interaction term in the logistic regression (‘GxS’) which maintains full sample size, and is implementable with standard software such as R/GenAbel (Aulchenko et al. 2007) and Plink (C. C. Chang et al. 2015). Despite the usability and power of these programs, principles of statistical genetics should be thoroughly understood prior to use. There has been extensive work on statistical methods for gene-by-environment interactions in GWAS, which can be applied to GxS (Gauderman et al. 2013). The GxS model does not appear to have been applied to any of the very large GWAS collections, possibly because of complications in the meta-analysis. Another potential issue with GxS is the potential to identify statistically-significant but biologically-unintuitive results. For example, a negligible difference in allele frequency between the sexes within the disease group, will generate a significant result if there is a similar difference in the opposite direction within the control group. Nevertheless, in a multi-collection GWAS, performing the GxS test for each subgroup, then performing the meta-analysis on these statistics may increase power over separate-sex meta-analysis.

The potential for false positives in GxS tests, created by an uneven sample size in each of the four groups, i.e. male and female for both cases and controls, enforces the need for large and carefully-selected control samples. Given the large number of different variables and co-factors in human populations (compared to controlled laboratory and agricultural studies), mixed models are being increasingly used in GWAS, and have good potential to investigate sex-dependent effects (Hoffman, Mezey, and Schadt 2014). Finally, it is now possible to conduct association tests across the X-chromosome that incorporates male hemizygosity and X-inactivation using the XWAS software package (Gao et al. 2015). This method has found new X-linked genes for auto-immune disorders, some of which exert sex-specific risk (D. Chang et al. 2014).

## Sex-dependent genetic effects by disease class

### Neurology and psychiatry

Although large GWAS have been performed on common neurological and psychiatric disorders, such as Parkinson's Disease, epilepsy, schizophrenia and bipolar disorder, there no robust reports of sex-dependent effects. In Alzheimer’s disease, the APOE gene, e4 risk allele exhibits an earlier onset in men but a greater overall risk in women, notably in heterozygous women carrying one APOE-ϵ4 allele (Riedel, Thompson, and Brinton 2016). The biochemical origins of this are expected to be in the lowering of bioenergetic rate during menopause, which creates a uniquely-female risk profile (Riedel, Thompson, and Brinton 2016). Within the other disorders, there are subtle sex-differences reported in prevalence, drug metabolism and clinical presentation, suggesting some influence of sex differences in pathogenesis and potentially genetics. (Also see Figure 1, Unipolar depression, with increased disability-adjusted life years in women globally.)

Autism typically has a 3:1 male-bias in prevalence, and is predominated by high-penetrance *de novo* mutations. These do not occur in male-biased genes, although gene expression in specific brain regions exhibits increased activity of normally-male biased genes (Werling, Parikshak, and Geschwind 2016). This provides strong evidence that hyper-masculinisation of certain regions of the brain, in both sexes, forms part of the aetiology of autism. In female autism cases, maternally-inherited genomic structural variation is a more common cause, leading to speculation that female embryos are more tolerant to this kind of genetic variation, but with the disadvantage of increased risk of disorders such as autism (Desachy et al. 2015). These examples for autism concern the impact of rare variants on disease, and are not under the same evolutionary forces as the high-frequency, low-risk variants which control many quantitative traits and common disease. Nevertheless, it constitutes a sex difference in genetic architecture, which is likely to originate from the differing pressures of natural selection on male and female genomes.

### Auto-immunity

These disorders classically show increased prevalence in females e.g. rheumatoid arthritis and osteoarthritis (from Figure 1 and see (Rubtsova, Marrack, and Rubtsov 2015), but there are few reports from high-powered GWAS of common immune disorders into the investigation or discovery of sex-dependent genetic effects. Application of the ‘XWAS method’ has identified X-linked sex-specific genetic effects on inflammatory bowel disease at genes C1-GALT1-C1, CENPI and MCF2 (D. Chang et al. 2014).

For asthma, a male-only risk effect has been observed at the ZPBP2 gene (Naumova et al. 2013). Although this result awaits replication in a larger population sample, there exists complimentary evidence: Firstly, common genetic variation at the same gene has recently been implicated in other common immune disorders such as systemic lupus erythematosus (Lessard et al. 2012) and ankylosing spondylitis (Qiu et al. 2013), albeit in a non sex-dependent manner. Secondly, the suppression of ZPBP2 activity by epigenetic methylation is greater in females than it is in males, providing a biochemical explanation for why only one sex is affected by the variant (Naumova et al. 2013), pending further investigation of tissue-specific methylation profiles. The reason why ZPBP2 is expression is not suppressed in males is possibly because it has an important role in production of viable sperm, but with no known phenotype in females, as determined from gene knock-out studies in mice (Y.-N. Lin et al. 2007).

### Metabolic: Cardiac, body-shape and type II diabetes

Robust sex-specific associations for cardiac-related quantitative traits such as thyroid hormone activity, waist-hip ratio, uric acid concentration, and triglycercides all of which are bio-markers for cardiac disease, have previously been reviewed (Gilks, Abbott, and Morrow 2014). For type II diabetes, two sex-dependent risk loci have also been identified from a GWAS meta-analysis of 149,000 participants, at CCND2 (male odds ratio 1.08–1.16) and at GIPR (female odds ratio 1.06–1.14) (Andrew P Morris et al. 2012).

Body shape, and density (as various metrics), is biologically-linked to cardiac disease and type II diabetes; It is simple and accurate to measure, which allows collection of large numbers of participants for high-powered studies. Recently, there have been four large GWAS meta-analyses of measurements of adiposity and body shape, which have generated many sex-dependent genetic effects for follow-up (n=7,44,20,4) (Randall et al. 2013; Winkler et al. 2015; Shungin et al. 2015; Lu et al. 2016). The trait most commonly studied was waist-hip ratio, adjusted by body-mass index (WHR_adjBMI_) which is a strong predictor of mortality caused by excess adiposity, often via heart disease and type II diabetes (Pischon et al. 2008). Sex-dependent loci which were identified in at least one of the other three studies include LYPLAL1, COBLL1/GRB14, VEGFA, and ADAMTS9, all of which have female-specific effects on WHR_a_djBMi (Figure 2). One study reported 11 out of 44 sex-dependent loci had significant opposing directions of effect between the two sexes (Winkler et al. 2015). Although these effects were not directly reproduced in the other studies, significant effects at some of the same genes were found, but in one sex only. Several genes now implicated in regulation of WHR_adjBMI_ in women, encode proteins that interact with one another. For example, PPARγ and RXRα proteins (Winkler et al. 2015) bind to form Adipocyte-specific transcription factor 6, a known master-regulator of adipocyte gene expression (Tontonoz et al. 1994). In another example, the transcription factors HoxC13 and MEIS1 bind *in vivo,* with complex tissue-dependent activity, and phenotypes relating to leukaemia, prostate cancer, hair, and limb development (Adamaki et al. 2015; Z. Lin et al. 2012)

Of the four GWAS studies, that by Shungin and colleagues (Shungin et al. 2015) is the most comprehensive and rigorous, although many results from the other studies are likely to be biologically meaningful. This study identified 49 loci effecting WHR_adjBMI_, of which 19 were female-specific (38%), and 1 was male-specific. The distribution of effect sizes at significant genes, suggests that a subset female-specific loci have greater effect sizes than loci that effect both sexes equally (Figure 2). Overall, these results tentatively suggest that female-specific genetic effects are likely to make up a large proportion of additive heritability for WHR_adjBMI_, which in-turn is linked to adiposity, heart disease and type II diabetes. However more information is needed on what the biological basis of WHR_adjBMI_ is, and whether it is correlated with mortality equally in both males and females.

**Figure 2.**
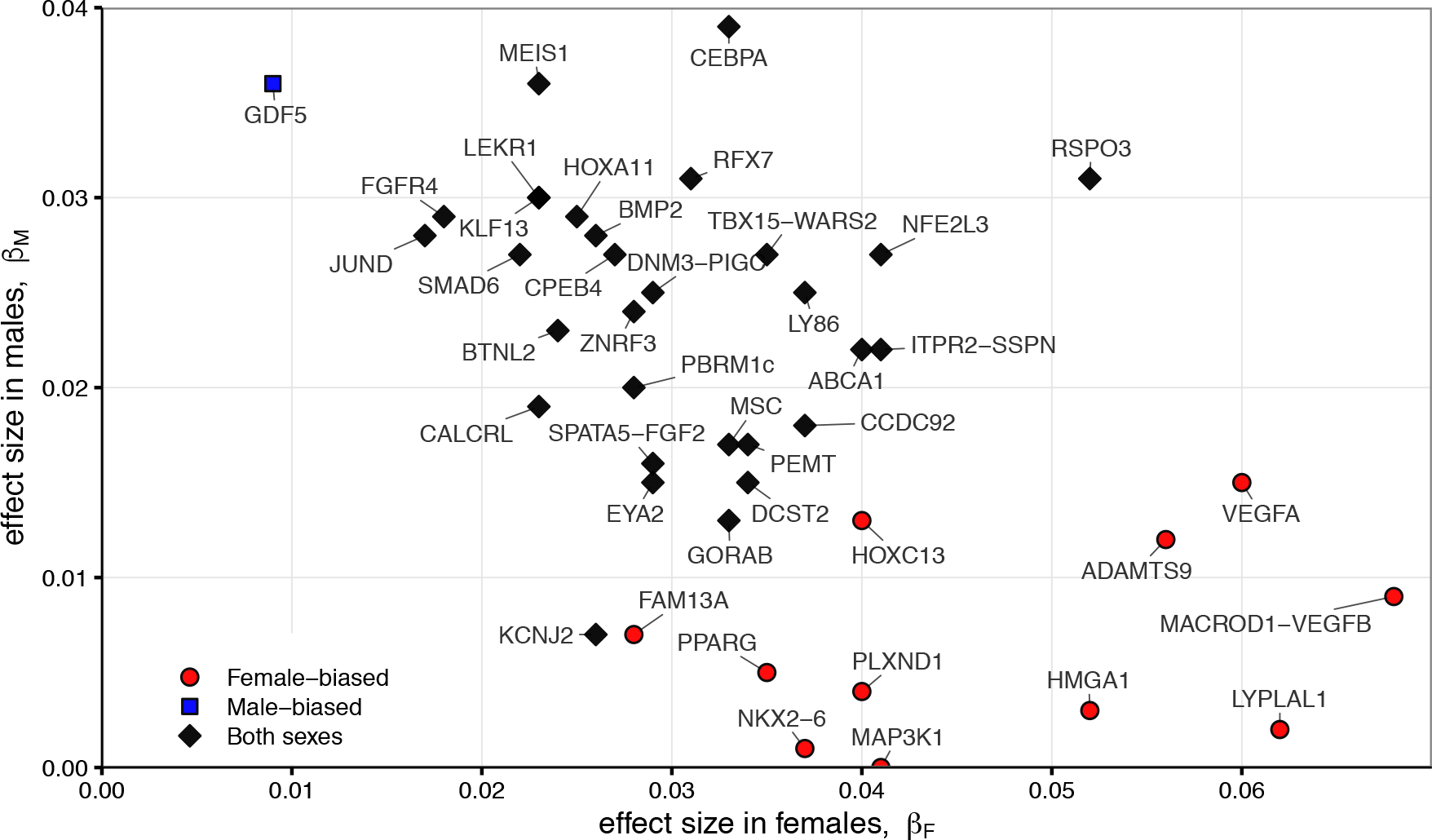
Male and female effect sizes for genes influencing waist-hip ratio (adjusted by body-mass index) in a study of 224,459 participants (Shungin et al. 2015). Loci which were calculated to have significant sex-dependent effects are coloured, red for females, blue for males.

### Cancer

Despite profound sex differences in prevalence in a variety of cancer types, and large GWAS collections, few robust sex-dependent genetic effects have been identified. Data presented in Figure 1 (top row) shows that global disability-adjusted life-years due to cancers of the liver, oesophagus and lung are all greater in males (WHO 2016), which is perhaps attributable to the generally higher rates of alcohol and tobacco use in this sex. Nevertheless, the only evidence for any sex-dependent genetic effect (other than those restricted by anatomical differences), is in males, whereby a missense variant in the aurora kinase 0 gene (AURKB) was shown to increase micro-nucleus formation (a cellular marker of carcinogenesis) in males only (McIntyre et al. 2016).

## Discussion

Sex-dependent risk loci are expected for common diseases because of sex-dependent expression quantitative trait-loci (eQTL), whereby expression level of a gene is determined by the combination of sex and genotype. In cell culture sex-dependent eQTLs constitute 15% of genes, and it is interesting to note that many genes with no difference in expression level between the sexes, can still have sequence variation which acts in a sex-dependent way on expression (Dimas et al. 2012; Werling, Parikshak, and Geschwind 2016). One biochemical possibility for this is that different transcription factors transcribe the gene at the same rate in each sex, but have different binding sites on the DNA, so only one of which will be effected by a short polymorphism. Sexually dimorphic gene expression varies dynamically by tissue type, age and chromosome (Kang et al. 2011; Lowe et al. 2015). For example, early-stage blastocysts have high levels of sexual dimorphism in gene expression presumably originating from the sex-determination cascade (Lowe et al. 2015). Sexually dimorphic gene expression in three regions of the neocortex is most common prior to birth, with up to 60 autosomal genes being expressed only in males, which by adulthood has dwindled to a few female-biased X-linked genes (Kang et al. 2011). In the liver, most of the dimorphism is in adult life but a small subset of genes here are dimorphic throughout all ages of life (Lowe et al. 2015). Furthermore, sexually dimorphic expression pre-implantation, is driven by sex-chromosome based transcription, whilst later development is characterised by sexually dimorphic autosomal transcription. These examples demonstrate how sexually dimorphic gene expression can be categorised, and how univariate analysis might be misleading.

The evolution of sex-dependent genetic effects on disease risk has been postulated to arise from intra-locus sexual conflict, but direct evidence is limited (Gilks, Abbott, and Morrow 2014). In the example of ZPBP2 and asthma, the increased gene activity and risk in males is likely to be maintained in the population because of the fundamental need for sperm production. For genomic structural variation in autism, if female embryos are indeed more resistant, this increases the female birth ratio but at, the price of reduced cognition (Desachy et al. 2015). Variants in RXRα increase waist-height ratio in females, and high expression levels correlated increased rates premature birth, which is inevitably associated with increase infant mortality. This might indicate balancing selection between nutrient metabolism and storage in the mother, and investment in the health of the offspring. Overall, the evidence hints that processes such as sex-specific pleiotropy and inter-locus sexual conflict might provide evolutionary explanations for the observed sex-dependent disease risk genes.

## Summary

The identification of sex-dependent loci for adiposity, type II diabetes and auto-immune disorders, in the largest sample collections, should provide incentive to screen other diseases. Furthermore, there is now evidence that a sub-group of sex-dependent risk loci have greater effect sizes than conventional, main-effects loci (e.g. for WHR_adjBMI_: ADAMST9, VEGFA, VEGFB, LYPLAL1, COBLL1, HMGA1). The greater effect size might originate from nonsynonymous (protein-altering) alleles, or from, more conserved pathways. Indeed, sexual dimorphism in gene expression can be classified by tissue type, chromosomal origin and varies dynamically with development and aging. Furthermore, evolutionary theory predicts that sex-dependent disease risk alleles will have greater effect sizes because they are only selected against in one sex (Morrow and Connallon, 2013). One cause for concern is the use of analysis methods which do not directly make a test of genotype-by-sex interaction, and thus remain under-powered.

There has been great public investment into GWAS meta-analyses, publications and results are becoming progressively more accessible, so release the computational code and logs detailing precisely how the analyses were performed might be of benefit to the rest of the research community. This review highlights the need, not only for more research into sex-dependent genetic effects, but also for continued collection of large sample sizes for GWAS, and a for comprehensive study of gene expression across all ages, sexes and tissue types, both of which are huge projects requiring massive collaboration, astute management, and fearless science. Smaller-scale studies are warranted too, simulations of experimental design and analysis methods for detecting sex dependent genetic effects, and investigation of the forces of evolution that provide deep-rooted explanations for the everyday problems of heritable diseases.

## Acknowledgements

The author is supported by the European Research Council, Starting Grant #280632 awarded to Dr Ted Morrow (University of Sussex), and acknowledges further stimulating discussions with Drs Alison Merikangas (University of Pennsylvania), Fiona Ingleby, and Tanya Pennell (both University of Sussex).

## URLs for data, software and code

Figure 1 WHO data http://www.who.int/healthinfo/global_burden_disease/estimates/en/index2.html

Figure 1 code http://dx.doi.org/10.5281/zenodo.53742

Figure 2 data and code http://dx.doi.org/10.5281/zenodo.57582

Plink/1.9 https://www.cog-genomics.org/plink2

GenABEL http://www.genabel.org/

XWAS http://keinanlab.cb.bscb.cornell.edu/content/xwas

## Further reading

- Kukurba KR, Parsana P, Balliu B, et al (2016) Impact of the X Chromosome and sex on regulatory variation. Genome Res. 26(6):768-77. doi: 10.1101/gr.197897.115.
- Rigby N, Kulathinal RJ (2015) Genetic Architecture of Sexual Dimorphism in Humans. J Cell Physiol 230(10):2304-10. doi: 10.1002/jcp.24979.
- Brown EA, Ruvolo M, Sabeti PC (2013) Many ways to die, one way to arrive: how selection acts through pregnancy. Trends Genet. 29(10):585-92. doi: 10.1016/j.tig.2013.03.001
- Cahill L (2012) A half-truth is a whole lie: on the necessity of investigating sex influences on the brain. Endocrinology 153(6):2541-3. doi: 10.1210/en.2011-2167
Mank JE, Hosken DJ, Wedell N (2014) Conflict on the sex chromosomes: cause, effect, and complexity. Cold Spring Harb Perspect Biol. 6(12):a017715. doi: 10.1101/cshperspect.a017715.

